# Sculpting DNA-based synthetic cells through phase separation and phase-targeted activity

**DOI:** 10.1101/2023.03.17.533162

**Authors:** Layla Malouf, Diana A. Tanase, Giacomo Fabrini, Miguel Paez-Perez, Adrian Leathers, Michael J. Booth, Lorenzo Di Michele

## Abstract

Synthetic cells, like their biological counterparts, require internal compartments with distinct chemical and physical properties where different functionalities can be localised. Inspired by membrane-less compartmentalisation in biological cells, here we demonstrate how micro-phase separation can be used to engineer heterogeneous cell-like architectures with programmable morphology and compartment-targeted activity. The synthetic cells self-assemble from amphiphilic DNA nanostructures, producing core-shell condensates due to size-induced de-mixing. Lipid deposition and phase-selective etching are then used to generate a porous pseudo-membrane, a cytoplasm analogue, and membrane-less organelles. The synthetic cells can sustain RNA synthesis *via in vitro* transcription, leading to cytoplasm and pseudo-membrane expansion caused by an accumulation of the transcript. Our approach exemplifies how architectural and functional complexity can emerge from a limited number of distinct building blocks, if molecular-scale programmability, emergent biophysical phenomena, and biochemical activity are coupled to mimic those observed in live cells.

## Introduction

Though there is no agreed definition of cellular life,^1^ there is general consensus on the fundamental characteristics of living cells, which includes information processing, adaptability, growth and division, metabolism, and compartmentalisation.^2,3^ Bottom-up synthetic biology aims to create synthetic cells featuring a subset, or all, of these fundamental characteristics by combining elementary molecular building blocks.^2,4^ This radical approach, although regarded as more challenging compared to traditional top-down cell engineering, circumvents the complexities inherent to working with and modifying living cells.^1,5^ Particularly challenging in the context of synthetic cell engineering is the production of sufficiently complex and robust cell-like architectures containing multiple compartments that serve to localise, segregate, and regulate function and environment.^6,7^ These enclosures are often membrane-based, constructed from phospholipids and/or fatty acid vesicles,^8–12^ polymersomes,^13,14^ or proteinosomes, ^15,16^ but alternative membraneless architectures are emerging in the form of hydrogel capsules, coacervates, or synthetic condensates.^17–21^

Microfluidics is a typical strategy for producing synthetic cell scaffolds, being particularly effective at generating monodisperse and nested structures in small quantities.^22^ However, the microfluidic approach can be difficult to implement and scale, requiring bespoke, often complex chips and specialised equipment.^23^ Like bulk emulsification approaches, microfluidics often rely on water-oil mixtures to assemble stabilised droplets or giant vesicles, which adds challenges and steps in the workflow associated to handling and removal of the non-aqueous phase.^24–33^ It may instead be more desirable to use techniques which exploit simple physical principles such as phase separation and self-assembly while avoiding the use of specialised equipment and non-aqueous components altogether.

The low costs of DNA oligonucleotides, ^34^ in conjunction with the predictable thermodynamics and kinetics of their interactions,^35,36^ and the availability of computational design tools,^37,38^ makes DNA nanotechnology ^36,39,40^ highly attractive to engineer advanced biomimetic devices, including synthetic cells. The latter are often formed from the programmable condensation of DNA building blocks in aqueous environments, hence removing the complexities of microfluidics, emulsification, and other chemical manufacturing routes. Artful examples of DNA-based, cell-like devices have been reported, self-assembled from multiblock single-stranded DNA,^41,42^ or from branched DNA junctions known as nanostars.^18–20,43,44^ Among condensate-forming DNA motifs, amphiphilic nanostructures, obtained by labelling DNA junctions with hydrophobic moieties, have shown significant potential owing to the stability of the self-assembled phases, their programmable nanostructure, and the possibility of engineering localised functionality and response to various environmental stimuli.^45–53^

While DNA-based synthetic cell implementations hold substantial promise due to their facile preparation, robustness, and algorithmic programmability, routes towards generating architectures with sufficiently complex internal structure, akin to those achievable with microfluidic methods, are lacking. In this work, we take advantage of the programmable phase behaviour of amphiphilic DNA nanostructures alongside their ability to host stimuli-responsive elements and sustain enzymatic pathways, ^46–48,51^ to construct cell-like devices that display a complex internal architecture and spatially resolved activity. The DNA-based synthetic cells feature a porous lipid shell as a pseudo-membrane, and internal membrane-less organelles. These organelles can be engineered to exhibit different functionalities, from selective etching which creates a cytoplasm-like space within the lipid shell, to hosting the *in-situ* transcription of RNA aptamers by a polymerase, which also induces spatially localised morphological changes to the synthetic cells. Our results demonstrate that the synergy between biophysical and biochemical responses can produce structural and functional complexity akin to that observed in live cells, even in minimalist systems composed of a small number of molecular building blocks.

## Results and discussion

### Generating internal heterogeneity in DNA condensates with phase separation

Our synthetic cells are primarily constructed from C-stars — DNA junctions made amphiphilic through the addition of cholesterol moieties — whose structure is sketched in Fig. 1(a). Subject to a one-pot thermal annealing, single-component samples of four-arm C-stars have been shown to self-assemble into polyhedral crystalline aggregates^45,46^ or, as shown in Fig. 1(b), spherical, cell-size condensates. ^46^ The latter, despite their macroscopic appearance, can be either crystalline or amorphous.^46,51^ Aggregate morphology (whether polyhedral or spherical) and the degree of crystallinity have been found to depend on C-star design features, size, quenching rate, and ionic conditions. ^45–47^ The nano-porous structure of the condensates enables internal diffusion of oligonucleotides and proteins. ^51^

**Figure 1:**
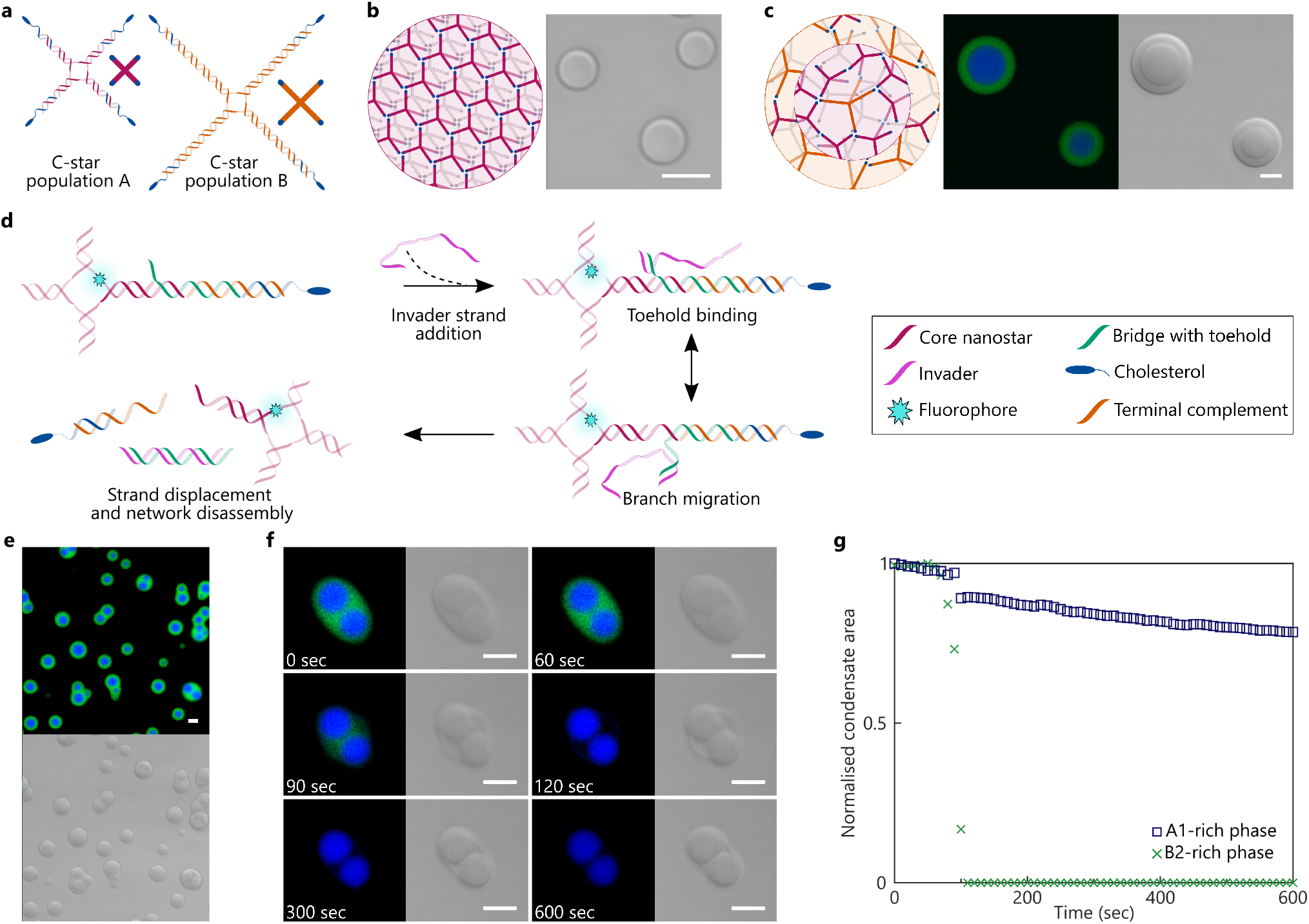
Size-induced phase separation leads to the emergence of core-shell DNA condensates and phase-targeted disassembly. **(a)** Schematic of two amphiphilic DNA nanostar (C-star) populations, A and B, with different arm lengths — A with 35 base-pairs (bp) and B with 50 bp arm length. The end of each arm is decorated with a cholesterol molecule.^45^ Simplified schematics adjacent. **(b)** Schematic (left) and bright-field micrograph (right) of C-star condensates comprised of C-star motif A1, with a 35 bp arm length. Condensates form upon annealing singlestranded C-star components from high temperature (see Methods)^45–47,52^ Within the condensates, C-stars interact through hydrophobic forces, with the cholesterol moieties likely co-localising in micelle-like regions. **(c)** Schematic (left), and confocal (middle) and bright-field (right) micrographs of binary, de-mixed C-star condensates comprised of C-star populations A1 (Cy5-labelled, blue) and B3 (fluorescein-labelled, green, 50 bp arm length). When coannealed, two different C-star populations self-assemble to form phase-separated condensates with distinct core-shell morphology, with the small-arm-length constructs localising primarily in the core, and the larger motifs favouring the shell. **(d)** Schematic of C-star disassembly driven by toehold-mediated strand displacement.^46^ An invader strand binds to the toehold and displaces a bridge strand, triggering condensate disassembly by dissociating the central C-star junction from cholesterol-DNA micelles. **(e)** Confocal and bright-field micrographs of another binary C-star condensate system, comprised of populations A1 and B2, both of which have been labelled with a fluorescent probe on their inner junction — Cy5 for population A1 (shown in blue, inner phase), and fluorescein for population B2 (shown in green, outer phase, arm length 50 bp). **(f)** Confocal and bright-field micrographs showing the targeted disassembly of the outer, B2-rich, phase of the binary condensate described in panel (e), exploiting toehold-mediated strand displacement as shown in panel (d). Timestamps mark time elapsed after adding the invader strand. **(g)** Disassembly of the condensate shown in panel (f) quantified by monitoring the cross-sectional area of the two phases measured with image segmentation. Abrupt disassembly is seen for the outer, B2-rich, phase. The A1-rich inner phase shows a small, sharp shrinkage when the outer phase disassembles, ascribed to the presence of B2 motifs in the A1-rich phase (incomplete de-mixing). The steady decrease in apparent area of the A1-rich phase is an artefact of photobleaching. All scale bars 10 μm.

In this study, we primarily consider two populations of four-armed C-stars — A and B — with different lengths of the double-stranded (ds) DNA arms: 35 base-pairs (bp) for type A and 50 bp for type B. Various designs for A-type (A1, A2) and B-type (B1, B2, B3) C-stars, were used, hosting different modifications. The oligonucleotide components of each motif and their sequences are listed in Supplementary Tables S1 and S2.

Figure 1(c) shows that condensates formed by A-B C-star mixtures (A1-B3 in this case) display two nested regions, a core enriched in the smaller construct A (identified through Cy5 labelling, shown in blue), surrounded by a spherical shell enriched in B (fluorescein-labelled, shown in green), as also confirmed by confocal Z-stacks in Supporting Video V1. Because A and B components can interact through identical cholesterol-cholesterol hydrophobic forces, phase separation is likely mediated by packing considerations driven by size mismatch, similar to de-mixing in binary mixtures of colloids of different size or colloid-polymer systems.^54–58^ Size-induced de-mixing may be enhanced by the tendency of C-stars to crystallise. ^45,46^ The core-shell morphology can be rationalised with interfacial energy considerations, with the Brich phase likely to have a lower surface energy for interaction with water due to its lower hydrophobic content (cholesterol/DNA molar ratio) compared to the A-rich phase.^59,60^

Size-induced phase separation is a simple but robust bottom-up approach for establishing addressable micro-environments within cell-like condensates, reminiscent of intracellular phase separation. ^61^ As we will explore later, individual building blocks can be modified with different responsive moieties, enabling spatial separation of functionalities. The robustness of size-induced phase separation means that the protocols developed for A-B systems are applicable to designs with slightly different arm lengths (C1, 48 bp and D1, 28 bp, see Supplementary Table S1). In addition, the formation of phase-separated binary condensates is straightforward and readily scalable, requiring a simple one pot thermal annealing (Methods).

### Targeted disassembly of C-star domains

As a first example of spatially-distributed functionality, we sought to enable the triggered disassembly of a selected C-star phase within the condensates, hence inducing a programmable, localised morphological response. This is preferable to the use of enzymatic approaches for condensate disassembly, such as deoxyribonucleases, which would degrade the DNA populations indiscriminately. ^62^ Selective network disassembly relies on toehold-mediated strand displacement, whereby an invader oligonucleotide displaces a DNA strand initially linking the sticky cholesterol moiety with the nanostar, as described in earlier work by Brady *et al.,*^46^ and shown in Fig. 1(d). Following disassembly, the targeted C-star motifs split into non-cholesterolised DNA stars and dispersed cholesterol-DNA micelles. ^46^

Figure 1(e) shows phase-separated binary condensates in which the larger, B-type, C-stars in the outer phase (fluorescein-labelled B2, shown in green) are modified to disassemble, while A-type C-stars accumulating in the core are non-responsive (Cy5-labelled A1, shown in blue). Condensate disassembly following the addition of the invader strand can be observed in Fig. 1(f) as loss of signal from the B2 channel. Figure 1(g) shows the time evolution of the cross-sectional area of the A-rich and B-rich phases for the condensate in panel (f), confirming the abrupt disassembly of the outer shell. Delay in the disassembly following invader addition (t = 0) results from its slow diffusion through the imaging chamber, while the progressive decrease in apparent area of the A-rich phase likely arises from the effect of photobleaching (visible in Fig. 1(f)) on the image segmentation pipeline (Methods). We also note a small, but sharp, decrease in the apparent area of the A-rich phase, simultaneous with B-phase disassembly. This shrinking response is likely a consequence of the disassembly of B-type motifs initially present in the A-rich phase, and the consequent relaxation of the remaining condensate. Incomplete demixing, namely the presence of B-type motifs in the A-rich phase (and *vice versa*), is expected in the binary condensates, and is further discussed and quantified below. Bright-field micrographs in Fig. 1(f) confirm the disassembly of most of the outer phase. However, low-density (low-contrast) material is left behind after the green phase has disappeared, which progressively coalesces with the A1-rich cores. Contrast enhancement of the fluorescence images, shown in Fig. S1, reveals a weak signal in the Cy5 channel, identifying the low density material as being composed of A-type C-stars initially present in the B-rich shell, consistent with incomplete de-mixing. Similar behaviour is observed in the disassembly of B3 C-stars in binary A1-B3 condensates, shown in Supporting Video V2.

### SUVs accumulate at the interface of C-star condensates, forming a lipid shell

Having shown how phase separation in multicomponent C-star condensates enables spatial engineering of functionality, we then proceeded to boost their biomimetic relevance by introducing a lipid layer. Small Unilamellar lipid Vesicles (SUVs, 100 nm in diameter, as confirmed by dynamic light scattering shown in Fig. S2), prepared from the zwitterionic lipid DOPC and stained with fluorescent Texas Red DHPE (0.8 % molar ratio), were introduced into samples of pre-formed, single component condensates (Methods). As sketched in Fig. 2(a), we observed adhesion of SUVs on the surface of the condensates, which appear to assemble into a continuous layer in confocal micrographs (Fig. 2(a), bottom right). We note here that the condensates have adopted a polyhedral morphology, reflecting the BCC crystalline phase formed by C-stars with this design (A1).^46^ As sketched in Fig. 2(a) (right), SUV adhesion is most likely mediated by the insertion of the C-star cholesterol moieties within the phospholipid bilayer - a known effect exploited for membrane functionalisation with DNA nanostructures. ^63–68^

**Figure 2:**
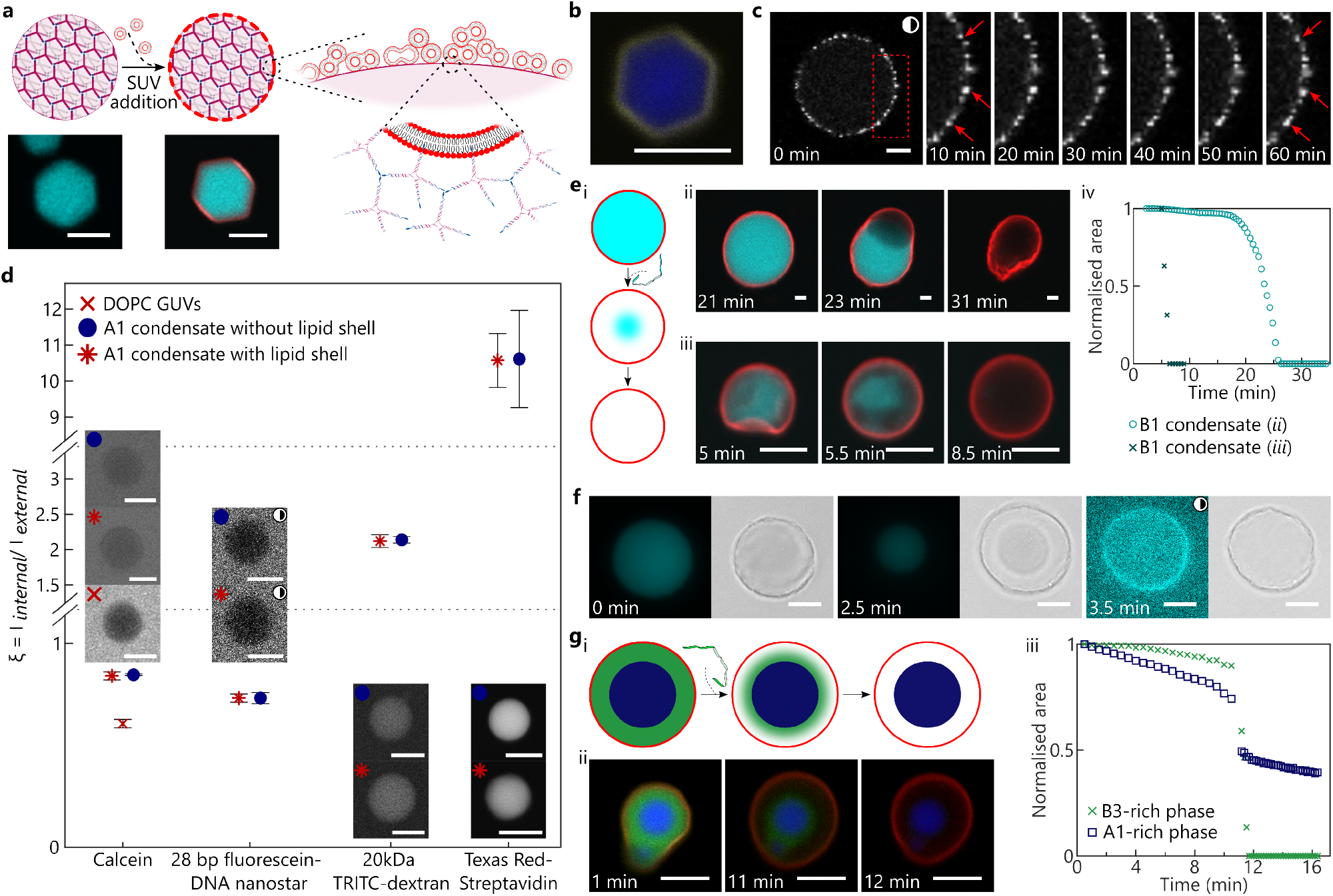
Liposome deposition leads to the formation of a porous lipid shell which remains stable after condensate etching. **(a)** Top: schematic of the lipid shell formed from the deposition of DOPC Small Unilamellar Vesicles (SUVs) on C-star condensates. Inset: proposed microstructure. Bottom: confocal micrograph of the Cy5-labelled A1 condensate (cyan) without (left) and with (right) a lipid shell labelled with Texas Red DHPE (red). **(b)** Confocal micrograph of a Cy5-labelled A1 C-star condensate (blue) with a lipid shell formed by calcein-loaded SUVs. Calcein fluorescent signal (yellow) demonstrates content retention in the adhered SUVs. **(c)** Confocal images showing the adhesion and lack of diffusion of SUVs on the surface of a C1 C-star condensate (arm length 48 bp). One in 800 SUVs are labelled with Texas Red DHPE (white), the remainder are non-fluorescent. Timestamps mark the time elapsed after acquiring the first image. **(d)** Partitioning of fluorescent molecular probes into A1 C-star condensates with and without a lipid shell, gauged using the ratio of probe fluorescence intensity inside (*I*_internal_) and outside the condensate (*I*_external_), measured from confocal micrographs (examples in insets). Symbols mark the mean of three independent repeats in which a median of 12 condensates were sampled, and error bars show the standard error. Data shows that the lipid shell is permeable to the tested probes, and that dextran and streptavidin preferentially accumulate within the condensates due the hydrophobic nature of TRITC and Texas Red. For calcein, *ξ* is also estimated for a sample of DOPC Giant Unilamellar Vesicles (GUVs), nominally impermeable to the dye. **(e)** Schematic (*i*) and confocal micrographs (*ii*) and (*iii*) of toehold-mediated strand displacement (TMSD) triggering disassembly of fluorescein-labelled B1 C-star condensates (cyan) encased in a Texas Red DHPE-labelled lipid shell (red). Timestamps mark time elapsed after invader strand addition. (*iv*) Disassembly of the condensates shown in panels (e)ii and iii quantified by monitoring the cross-sectional area of the condensates with image segmentation. Delays in onset of disassembly are a consequence of invader strand diffusion. **(f)** Epifluorescence and bright-field micrographs of TMSD disassembly of a fluorescein-labelled B1 condensate (cyan) encased in a non-fluorescent lipid shell. Timestamps mark time elapsed after adding the invader strand. After complete disassembly, at 3.5 minutes, enhancing the contrast of the cyan channel shows a fluorescent ring colocalised with the lipid shell, which remains after disassembly. **(g)** Schematic (i) and confocal micrographs (ii) showing TMSD disassembly of B3 C-stars in a binary condensate comprised of C-star motifs A1 (Cy5-labelled, blue) and B3 (fluorescein-labelled, green) encased in a lipid shell (Texas Red DHPE-labelled, red). (*iii*) Disassembly of the condensate shown in panel (g)ii quantified by monitoring the cross-sectional area of the two phases measured with image segmentation. As in Fig. 1(g), we observe abrupt disassembly of the outer, B3-rich, phase, here accompanied by a slight shrinkage of the A1-rich inner phase. All scale bars 10 μm, with the exception of 20 μm for panel (c). For images marked with a half-shaded circle, contrast enhancement has been applied by linear rescaling of the pixel intensity to aid visualisation.

In a related system, Walczak *et al.* reported the adhesion of small C-star aggregates onto the surface of cell-size Giant Unilamellar Vesicles (GUVs),^48,53^ observing that C-star particles are able to permeabilise liposomes and cause their rupture. To assess whether similar destabilisation occurs for SUVs depositing onto cell-size C-star condensates, we encapsulated calcein inside SUVs lacking phospholipid fluorescent labelling. Figure 2(b) shows that the calcein signal (shown in yellow) remains localised on the condensate surface, confirming that (at least some of) the SUVs are not disrupted nor rendered leaky. We then prepared lipid shells in which only a small fraction of the SUVs (1 in 800) were doped with Texas Red DHPE, while the remainder consisted entirely of unlabelled lipids. As shown in Fig. 2(c), we observed a speckled pattern on the lipid layer, whose persistence over time demonstrates that neighbouring SUVs are unable to exchange fluorescent lipids and are thus not undergoing significant fusion. Monitoring the speckle arrangement over time shows no Brownian motion, indicating that the SUVs remain effectively static over experimentally-relevant timescales. Taken together, the absence of substantial leakiness and lack of lipid exchange or diffusion hints at a morphology of the lipid layer as depicted in Fig. 2(a), in which the SUVs mostly retain their identity rather than reconfiguring into a continuous bilayer.

It is thus expected that the lipid layer would have significant porosity. This hypothesis is tested in Fig.2(d), where we used confocal microscopy to measure the permeability of the lipid layer. C-star condensates formed from the A1 motif, either with or without a lipid shell, were soaked in solutions of various fluorescent probes. The ratio (*ξ*) of fluorescent intensities recorded within (*I_internal_*) and outside (*I_external_*) the condensates was used as a proxy to determine whether the lipid shells are permeable to these probes (see Methods). ^46^ For all tested probes, which included calcein, a non-cholesterolised fluorescein-labelled DNA nanostar with 28 bp arm length, 20 kDa tetramethylrhodamine isothiocyanate (TRITC)-labelled dextran, and Texas Red-labelled streptavidin, we observe that *ξ* is unaffected by the lipid shell. This evidence confirms that the shell possesses pores sufficiently large to allow for diffusion of all tested probes. As a comparison, we present data for penetration of calcein into electroformed GUVs, which should be largely impermeable to the dye. As expected, *ξ* is significantly lower for GUVs compared to condensates, although greater than zero due to out-of-plane fluorescence signals. We also note that dextran and streptavidin accumulate within the condensates, driven by favourable hydrophobic interactions between the cholesterol moieties and the hydrophobic, rhodamine-derived fluorophores they host. ^46^ A similar test shows that shell porosity is not reduced with longer condensate-SUV incubation times (Fig. S3).

### Lipid shells remain stable after condensate disassembly

The porosity of the lipid shell allows rapid exchange of material, which we can exploit to externally trigger the disassembly response discussed earlier, through the addition of an invader strand. In Fig. 2(e) we observe condensates of responsive B1 C-stars dissolve from within the lipid shell, confirming that the latter is sufficiently porous to first allow inward diffusion of the invader, and later the escape of the disassembled DNA fragments (the largest of which is a non-cholesterolised DNA star with 12 bp arm length). Surprisingly, we note that, in the vast majority of cases, the lipid shells remain stable after condensate disassembly is complete. Therefore, despite the previously noted evidence that SUVs largely retain structural identity (Fig. 2), a mechanism must exists through which the SUVs remain physically connected, as we will discuss further.

Lipid-coated condensates display diverse responses upon disassembly, two examples of which are shown in Fig. 2(e). In panel (e) ii, the condensate disassembles asymmetrically, and a “bubble” is formed which deforms the lipid shell. In panel (e) iii, disassembly occurs more symmetrically but, as before, an expansion and smoothing of the lipid shell is observed. Though the behaviours differ, both examples suggest an initial build-up in osmotic pressure within the lipid shell due to the release of DNA nanostructures. ^20^ At later times, the pressure is released as the DNA diffuses out through the pores in the shell, evident in the absence of significant DNA-associated fluorescent signal remaining within the shells. Figure 2(e) *iv* shows the time-dependent cross-sectional area of the DNA condensates, confirming abrupt disassembly upon invader addition for the examples in Fig. 2(e) *ii* and iii. The different onset times noted for the two condensates is a consequence of the delayed diffusion of the invader in the imaging chamber (Methods). Disassembly of lipid-coated DNA condensates can also be performed by non-specific means, namely using DNase I. This is demonstrated in Fig. S4, where the lipid shell is also observed to persist once disassembly of a C-star condensate (population D1, arm length 28 bp) is completed.

Figure 2(f) shows TMSD disassembly of a B1 condensate coated in a non-fluorescent lipid shell. As expected, the fluorescent signal from the DNA in the condensate disappears as disassembly progresses. However, on enhancing image contrast post-etching, we note a faint signal co-localised with the lipid shell (Fig. 2(f), right). Supporting Video V3 shows a confocal z-stack of a Texas Red-labelled lipid shell remaining after disassembly of a fluorescein-labelled B1 condensate — with the contrast of the fluorescent channels enhanced, the same co-localisation of the DNA and lipid fluorescent signals is noted. We argue that a small amount of DNA nanostuctures may fail to disassemble in the region immediately in contact with the lipid shell, or in the gaps between the SUVs, possibly due to poor accessibility of the toehold domain. This protective effect of the lipids may be reminiscent of the one observed by Julin *et al.,* who found that cationic lipid-coated DNA origami were significantly more resistant to DNase I digestion than non-coated origami. ^69^

It is thus reasonable to hypothesise that residual amphiphilic DNA material may play a role in conferring stability to the lipid shell following condensate disassembly, by cross-linking neighboring SUVs. DNA-mediated SUV crosslinking would be compatible with the observations reported in Fig. 2(b) and (c) concerning the lack of content leakage and lack of significant lipid exchange between the SUVs in the shell. Nonetheless, other DNA-independent mechanisms may still play an important role, such as membrane adhesion or SUV hemifusion, encouraged by the presence of amphiphiles. ^70,71^

The ability to create semi-permeable lipid shells, combined with that of building condensates with distinct regions hosting addressable functionality, offers a route to construct architectures which resemble a prototypical biological cell. As demonstrated in Fig. 2(g), this can be achieved by forming a lipid shell around a binary A1-B3 condensate, in which the B3-rich outer phase can be disassembled with TMSD. Upon disassembly, a gap, reminiscent of a cytoplasm, is formed between the lipid shell and the A1-rich “organelles”. In Fig. S5, we observe a similar gap formed from the disassembly of the B3 motif in A1-B3 binary condensates enveloped in a lipid shell comprising SUVs encapsulating calcein. Here, we note the calcein signal persisting, indicating a lack of significant content leakage. As demonstrated in Fig. 1(g), disassembly can be quantified by monitoring the cross-sectional area of the core and shell regions. Consistently, in Fig. 2(g) *iii*, we note a sharp, complete disassembly transition for the B-rich phase and a simultaneous shrinkage of the A-rich core.

### In-situ transcription triggers morphological changes in DNA-based synthetic cells

In a recent contribution, Leathers *et al.* showed that C-stars can be modified with DNA templates, which are transcribed by T7 RNA polymerase to produce RNA aptamers *in situ*, thus allowing the DNA-based synthetic cells to produce new nucleic acid building blocks. ^51^ Here we seek to combine these transcription capabilities with the more complex cell-like architectures we present, which differ from the implementation of Leathers *et al.* by featuring regions with distinct physical characteristics and composition.

We consider binary systems of C-stars that differ both in size and functionality, sketched in Fig. 3(a). One population is composed of B1 C-stars (50 bp arm length) which host a modification enabling disassembly through TMSD, as described previously. The second population comprises A2 C-stars (35 bp arm length), which are modified so that an overhang on one of the arms connects to a transcribable ssDNA Template (T) through a ssDNA Bridge (B) strand — see inset in Fig. 3(a). The Template codes for an extended and brighter version of the Broccoli RNA light-up aptamer, ^51^ which binds to and induces fluorescence in DFHBI, ^72^ while the Bridge contributes to forming the doublestranded promoter region required to initiate transcription by T7 RNA polymerase. ^51,73^

**Figure 3:**
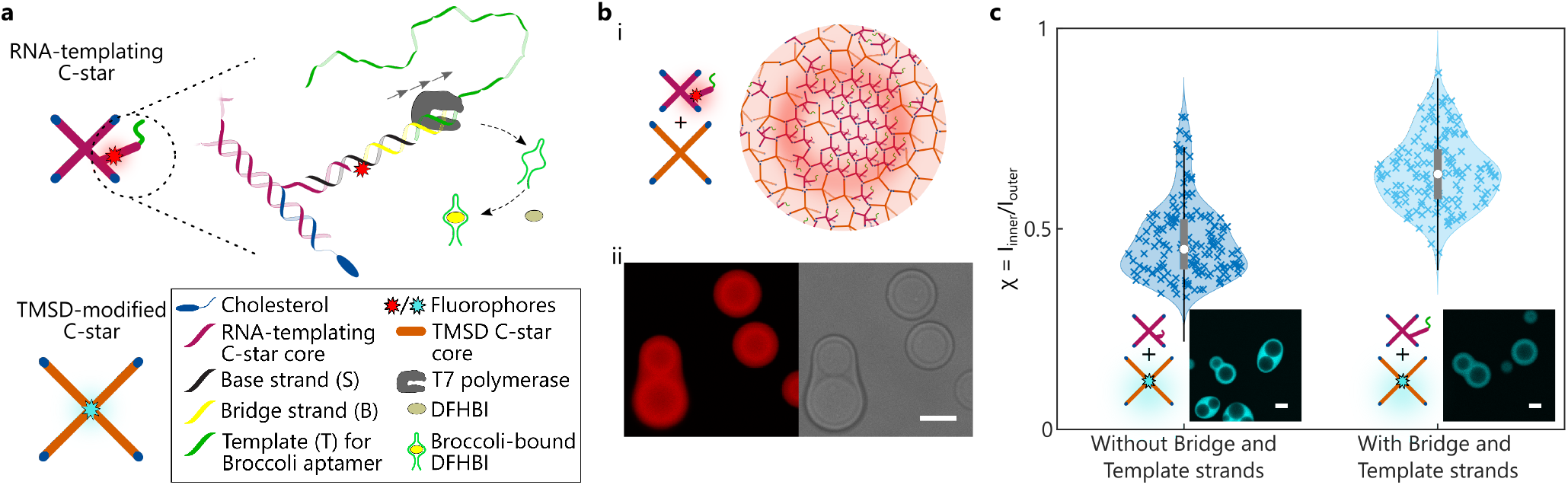
Modification of the DNA building blocks with transcribable constructs influence phase separation. **(a)** Simplified schematics of two C-star populations, one modified to introduce a DNA template, transcribable by T7 RNA polymerase,^51^ and another modified for TMSD-disassembly. Inset shows the ssDNA Base (S) and Bridge (B) strands linking the transcribable ssDNA Template (T) to the core nanostar. The Bridge may be labelled with Alexa Fluor 647, and also contains a DNA sequence, which, when hybridised with the complementary sequence on the Template strand, comprises the double-stranded T7 promoter sequence marking the transcription start site. The Template codes for an extended version^51^ of the RNA Broccoli aptamer, which binds to and induces fluorescence in DFHBI.^72^ **(b)** Schematics (*i*) and confocal and bright-field micrographs (*ii*) of a binary condensate comprised of both the RNA-templating C-star A2 and TMSD-modified C-star B1, where the former has been labelled with a fluorescent probe on the Bridge strand, showing the non-uniform distribution of the fluorescent signal from the Bridge strand. **(c)** Ratio *χ* of fluorescence intensity of the inner (*I_inner_*) and outer (*I_outer_*) phases of a binary condensate comprised of A2 and B1 C-stars, where the B1 motif is labelled with fluorescein and the A2 C-stars either contain all component strands or exclude B and T strands. Image segmentation (see Methods) is used to extract fluorescence intensity of 164 A2-B1 condensates annealed without the BT duplex, and 149 condensates annealed with the duplex. Data shows that the fluorescein-labelled B1 C-stars partition more strongly into the outer phase of the binary condensate when the BT duplex is not present; *i.e.* phase mixing increases with the introduction of the BT duplex. Violin plots show the median (white circle) and interquartile range (grey rectangle) of the data, with whiskers reaching ±1.5 × interquartile range from the first and third quartiles, shading showing a kernel density estimate of the data, and crosses marking the individual data points. All scale bars 10 μm.

As expected, one-pot annealing of all ssDNA components produced phase separated condensates, visible in bright-field images (Fig. 3(b)ii), right). However, when labelling the Bridge strand with a fluorescent probe (Alexa Fluor 647), confocal images reveal a non-uniform distribution of the Bridge and Template (BT) duplex in the inner phase, with a greater concentration found at the outer interface of the A-rich core compared to its centre (Fig. 3(b)*ii*, left). We also note a visible signal from the Bridge in the outer shell, indicating a significant presence of fluorescently-labelled A2 motifs in the B-rich region. A microscopy-informed schematic of the internal morphology of the condensates is sketched in Fig. 3(b)*i*.

Control experiments with A2-only condensates, summarised in Fig. S6, reveal that the presence of the bulky BT duplex is, in itself, sufficient to produce size-induced phase separation, leading to the appearance of a template-enriched shell and a template-depleted core, as visible by comparing condensates lacking and including Bridge and Template strands (Fig. S6(a) and (b), respectively). We thus argue that the non-uniform template distribution seen in Fig. 3(b) for binary condensates could be a direct consequence of size-induced phase separation caused by the template.

Besides exhibiting a non-uniform distribution, due to its substantial contribution to the molecular weight of the A-type stars, the BT construct may also influence the degree of phase separation between A-type and B-type motifs in the binary condensates. To test this hypothesis, we prepared binary condensates comprising fluorescein-labelled B1 C-stars and unlabelled A2 C-stars, the latter of which either contained or did not contain the Bridge and Template strands. Confocal microscopy confirms that in both condensate types, the outer phase is enriched in fluorescein-labelled B1 C-stars (shown in cyan in Fig. 3(c)). Using image segmentation, we extracted the average B1 fluorescent signal in the inner A-rich phase (*I*_inner_) and in the outer B-rich phase (*I*_outer_), and computed the parameter *χ* = *I*_inner_/*I*_outer_. *χ* can be used as a proxy for the extent of A-B mixing in the condensates, taking values close to zero for complete de-mixing and close to 1 for complete mixing. A higher value of *χ* is found when BT constructs are present, indicating that the modifications hinder A-B phase separation, likely by making the steric encumbrance of A-type motifs closer to that of B-type motifs. It should also be noted that, even in the absence of BT, *χ* remains substantially larger than 0 (~ 0.45) indicating incomplete de-mixing and consistent with the observations made on the response of the system to disassembly of the outer phase (Fig. 1(g) and Fig. 2(g)).

Having characterised phase separation and BT-distribution in A2-B1 binary mixtures, we proceeded to create cell-like architectures able to sustain transcription, henceforth referred to as synthetic cells. To this end, the BT-containing A2-B1 binary condensates were coated in a lipid shell as discussed above, and washed multiple times to remove any free DNA and unattached SUVs (see Methods). The removal of freely-diffusing Broccoli-templating DNA was verified with transcription reactions run using the supernatants extracted after each wash, as shown in Fig. S7(a). The B1 motifs were then disassembled with TMSD, as sketched in Fig. 4(a). Epifluorescence images show a significant quantity of labelled RNA-templating A2 C-stars (red) in the B1-rich outer phase pre-etching, as expected given the hindering effect that the BT duplex has on A-B de-mixing, quantified in Fig. 3(c). During disassembly of the B1 C-stars, some of the BT-linked A2 motifs collapsed to form a low density mesh surrounding the core of the synthetic cells. Other constructs were able to detach and escape the lipid shells, as confirmed by control experiments in which RNA transcription reactions are carried out with the supernatants removed from the samples post-B1 etching, and summarised in Fig. S7(b). Despite the synthetic cells having been sufficiently washed pre-etching (Fig. S7(a)), the Broccoli signal from the supernatant was again substantial post-etch, indicating a significant leakage of the Broccoli-templating A2 constructs during removal of the outer phase (Fig. S7(b)). Bright-field micrographs in Fig. 4(b) also show the lipid shell shrinking and wrinkling during the disassembly, more substantially than for other, template-free, designs (Fig. 2(g) *ii*). As we discuss later, this response is likely a result of less complete de-mixing, where a significant quantity of A2-rich material is present in the outer phase and collapses onto the inner core after etching, which may pull on the lipid layer causing its contraction.

**Figure 4:**
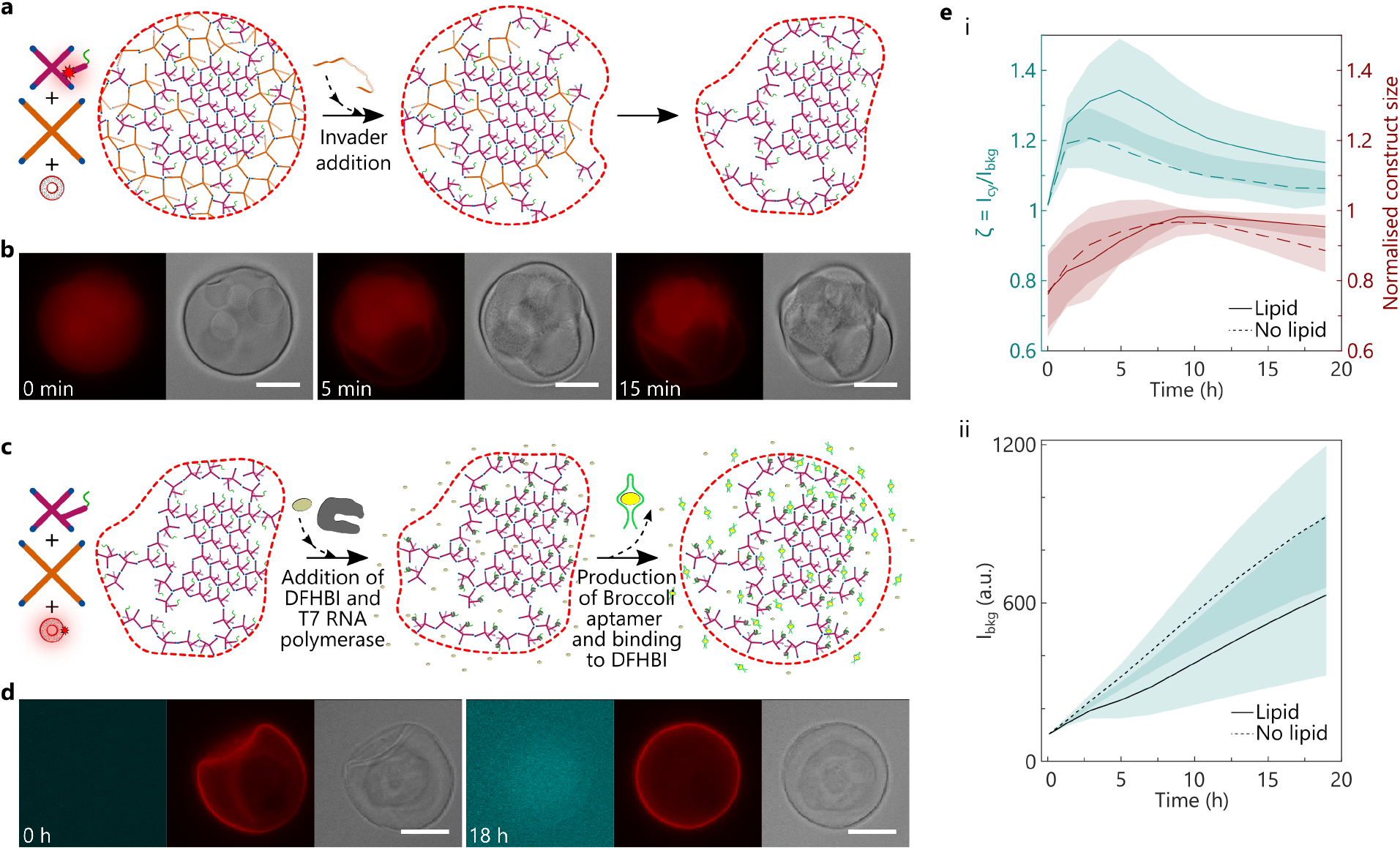
RNA synthesis induces morphological changes in synthetic cells. **(a)** Schematic showing the disassembly of B1 C-stars in a lipid-coated A2-B1 binary condensate (synthetic cell), triggered by addition of an invader strand inducing a TMSD reaction (Fig.1(d)). **(b)** Brightfield and epifluorescence micrographs showing the TMSD reaction sketched in (a). In the fluorescent images, the red signal marks the location of the Bridge strand labelled with Alexa Fluor 647 (red). Lipids and other DNA components are unlabelled. Note that etching of the B1-rich phase leads to shrinking of the synthetic cell. From the bright-field images, one can observe the re-structuring of the outer C-star phase following TMSD, which forms a mesh-like layer surrounding the A2-rich cores. Timestamps mark time elapsed after invader strand addition. **(c)** Schematic showing Template transcription and Broccoli synthesis in post TMSD synthetic cells where the lipid shell is labelled with a fluorescent probe. **(d)** Brightfield and epifluorescence micrographs of the process depicted in (c). The fluorescence signal from Broccoli·DFHBI, shown in cyan (left), increases over time, and is accompanied by swelling of the synthetic cell. The Texas Red-labelled lipid shell is shown in red (middle), and is observed to remain intact during transcription. Timestamps mark time elapsed after addition of T7 RNA polymerase and transcription mixture (see Methods). **(e)** Quantification of Broccoli synthesis and morphological changes for synthetic cells undergoing transcription, where the lipid shell is non-fluorescent and the Bridge strand is labelled with Alexa Fluor 647 (as in (a), (b) and Figs. S6 and S7, and Supporting Videos V8 - V13). (*i*) Left axis: Ratio between the fluorescence intensity of Broccoli·DFHBI sampled from epifluorescence micrographs in the region surrounding the A2-rich core (“cytoplasm”) of the synthetic cells, and in the sourrounding solution (*I_bkg_*). Right axis: Condensate size. (ii) Time-dependence of *I_bkg_*. Solid lines and dashed lines are relative to synthetic cells featuring or lacking a lipid shell, respectively. For both panels (e)i and ii, data is extracted through image segmentation (see Methods and Fig. S11) of 23 lipid-coated (solid line) condensates and 16 non-coated (dashed line) synthetic cells; with lines showing the mean and shaded regions indicating the mean ± the standard deviation of the data. All scale bars 20 μm.

Transcription of the etched and subsequently washed synthetic cells causes an expected increase in Broccoli fluorescent signal, and a distinct morphological response, both of which are sketched in Fig. 4(c) and shown in epifluorescence micrographs in Fig. 4(d). Broccoli fluorescence builds up within the A2-rich dense inner phase, within the cytoplasm-like region and low-density material left behind by B1 disassembly, and later in the medium surrounding the synthetic cells. More surprisingly, we note a dramatic change in the morphology of the disordered A2-rich mesh surrounding the core of the synthetic cells, which expands and inflates the synthetic cell to a size and shape akin to that observed prior to etching. Additional images and further examples of the transcription-induced synthetic-cell expansion can be found in Figs. S8-S10 and Supporting Videos V5-V13. Fluorescent labelling of the SUVs used in Fig. 4(d), Fig. S8 and Supporting Videos V5-V7 demonstrates that the lipid shell follows the expansion the synthetic cell pseudo-cytoplasm, which however also swells in the absence of lipids, as shown in Fig. S9 and Supporting Videos V11 - V13.

In order to rationalise this morphological response, we used image analysis to extract the outer dimensions of the synthetic cells and the intensity of the Broccoli fluorescence signal, measured both in the solution surrounding the synthetic cells (*I_bkg_*) and in their cytoplasm (*I_cy_*, see Fig. S11). The results of this analysis are presented in Fig. 4(e)*i* for synthetic cells prepared with the Alexa Fluor 647-labelled Bridge strand, with or without a (non-fluorescent) lipid shell. Regardless of the presence of a lipid shell, construct size increases as transcription progresses and reaches a maximum after approximately 8 hours, followed by a slight decrease in size which is less pronounced for the lipid-coated synthetic cells. The fluorescence intensity ratio *ζ* = *I_cy_/I_bkg_* should approximate the ratio of aptamer concentrations inside and outside the constructs. As might be expected, at early times, we see a pronounced growth of ζ from its initial value of ~ 1 due to localised Broccoli transcription within the synthetic cells. A maximum is then reached, followed by a steady decrease back towards ζ ~ 1, due to aptamer leakage and the consequent increase of *I_bkg_* (also notable in Figs. S9 and S10 and Supporting Videos V8-V13), combined with the progressive reduction in polymerase activity. A clear difference is observed between lipid-coated and non-coated constructs, with the former displaying a higher maximum ζ-value, which is also reached at later times. These differences indicate that, albeit permeable, the lipid shell is able to slow down outward diffusion of the Broccoli aptamer. This hypothesis is consistent with the absolute values measured for *I_bkg_*, which are higher for lipid-less constructs compared to the complete synthetic cells, as shown in Fig. 4(e) *ii*. The accumulation of the Broccoli aptamer within the synthetic cells (whether lipid-coated or not) hints at potential mechanism for the observed size increase, where a transient osmotic pressure build-up from the transcript causes swelling of the low-density A2-rich material remaining around the cores of the constructs following removal of the B1-rich shell. A similar response was observed by Saleh *et al.* during the enzymatic digestion of DNA hydrogel droplets, where the formation of internal cavities was ascribed to osmotic pressure from disassembled DNA fragments. ^20^

## Conclusion

In summary, we have presented a strategy for constructing cell-like architectures that mimic multiple characteristics of biological cells, including a porous lipid-based shell, a cytoplasm-like cavity, and dense membrane-less internal compartments. These biomimetic devices are produced *via* bulk self-assembly of amphiphilic DNA nanostructures that undergo phase separation, combined with liposome deposition and phase-specific etching, hence negating the need for complex microfluidics or emulsion-based methods. The membrane-less organelles can be modified to host a DNA template. This template can be transcribed to produce RNA, whose localised synthesis causes a swelling response in the cytoplasm and lipid shell of the synthetic cells, due to transient osmotic pressure build up. The ability to initially accumulate, and then progressively release nucleic acids of tailored sequence could make our solution a valuable starting point for the development of therapeutic agents, *e.g.* for vaccines^74^ and gene therapy. ^75^

Our proof-of-concept implementation dovetails the structural and dynamic control afforded by DNA nanotechnology, with amphiphile self-assembly, phase-separation phenomenology, and *in vitro* transcription, hence exemplifying how increasingly more complex architectures and responses can be engineered from the bottom-up, if complementary molecular tools are synergistically combined. The synthetic cell assembly strategy we outline is general, and can be systematically expanded by localising functional moieties other than the DNA templates in the co-existing phases, *e.g.* grafted enzymes ^76^ or nanoparticles. ^77^ Despite being used here only to form a semi-permeable shell, the liposomes could further be targeted with lipid-specific functional modification, such as membrane receptors, while their ability to retain contents could be exploited to encapsulate and conditionally release molecular cargoes. This modularity and design versatility further strengthens the applicative outlook of our solution, potentially unlocking the design of multi-functional therapeutic synthetic cells that combine nucleic acid synthesis with targeting abilities and the possibility of co-delivering small-molecule drugs and macromolecules encapsulated in the liposomes and/or embedded in the DNA matrix. ^46^

## Acknowledgement

LM, LDM, and DT acknowledge support from the European Research Council (ERC) under the Horizon 2020 Research and Innovation Programme (ERC-STG No 851667 – NANOCELL). GF acknowledges funding from the Department of Chemistry at Imperial College London. MPP acknowledges support from a UK Research and Innovation New Horizons Grant (EP/V048058/1) and an EPSRC Doctoral Prize Fellowship (EP/W524323). AL and LDM acknowledge acknowledge support from a Royal Society Research Grant for Research Fellows (RGF/R1/180043). LDM also acknowledges support from a Royal Society University Research Fellowship (UF160152). MJB is supported by a Royal Society University Research Fellowship (URF/R1/180172) and acknowledges funding from a Royal Society Enhancement Award (RGF/EA/181009) and an EPSRC New Investigator Award (EP/V030434/2). The authors acknowledge the Facility for Imaging by Light Microscopy (FILM) at Imperial College London and thank Stephen Rothery for his assistance at the facility. The authors thank Elisa Franco for useful feedback on the manuscript.

