## Supplementary material for "Sculpting DNA-based synthetic cells through phase separation and phase-targeted activity": SI

#### Materials

DNA oligonucleotides were sourced from Integrated DNA Technologies (IDT) and purified by the supplier using standard desalting for non-functionalised strands, reverse phase high-performance liquid chromatography (RP-HPLC) for fluorophore-labelled strands, and polyacrylamide gel (PAGE) purification for "A2.Core1" and the Broccoli templating strand ("A2.Template"). 99% Texas Red 1,2-dihexadecanoyl-sn-glycero-3-phosphoethanolamine (DHPE-TR) was purchased from Stratech. 1,2-dioleoyl-sn-glycero-3-phosphocholine (DOPC) was purchased from Avanti Polar Lipids. 3,5-Difluoro-4-hydroxybenzylidene imidazolidinone (DFHBI), Sephadex G50, and tetramethylrhodamine isothiocyanate–dextran (TRITC-dextran, M.W. 20 kDa) were purchased from Sigma-Aldrich. Calcein was purchased from

Sigma-Aldrich and 100 mM stock solutions were prepared in Milli-Q water with 1 M NaOH (VWR) to bring the solution to pH ~7. Texas Red-labelled streptavidin was purchased from ThermoFisher Scientific. DNase I and its buffer (RNase-free) were purchased from New England Biolabs. RNA aptamer transcription experiments were executed using the CellScript T7-FlashScribe Transcription Kit purchased from Cambio. Sucrose (RNase- and DNase-free) was purchased from VWR. NaCl and Tris-EDTA (TE, 100 × stock) were purchased from Sigma-Aldrich. TE buffer was diluted in Milli-Q water prior to use, to an end concentration of 10 mM Tris, 1 mM EDTA, pH ~8.0. All buffer solutions were filtered through 0.22 $\mu$m syringe filters (Millex), stored at 4 °C and used within three weeks of preparation.

### Methods

#### C-star design and condensate preparation

Novel DNA sequences were designed and analysed using NUPACK.[1] Most oligonucleotide strands were reconstituted from lyophilised state in TE, with the exception of "invader" strands which were reconstituted in 0.3 M NaCl in TE. Reconstituted DNA strands were stored at -20 °C until used. Sequences of DNA strands used in this work are listed in Tables S1 and S2. Rectangular glass capillary tubes, purchased from CM Scientific, were cleaned by ultrasonication in a solution of 1 % Hellmanex III (Hellma) in deionised (DI) water for 30 min at 40 °C and then rinsed thoroughly with DI water. The capillary tubes were then further ultrasonicated in 2-propanol (Sigma-Aldrich) at 40 °C for 15 to 30 minutes and dried under $N_2$ prior to use. Two sizes of capillary tubes were used: large, with dimensions 0.4 mm × 4mm × 50 mm, and small, with dimensions 0.2 mm × 4mm × 50 . Condensates were prepared to a final concentration of 5 $\mu$M by mixing stoichiometric ratios of the appropriate strands for each population in 0.3 M NaCl in TE buffer. See Supplementary Tables S1 and S2. For binary C-star condensates, equal volumes of each population mix were combined into one container. An appropriate volume of these mixtures were pipetted into the cleaned glass capillary tubes (~60 $\mu$L in large tubes, and ~30 $\mu$L in small tubes), capped at both ends with mineral oil, and then sealed to a glass coverslip using Araldite Rapid 2-component epoxy glue. Samples were annealed using a Bio-Rad thermal cycler, and two annealing protocols were typically used:

- **Short anneal**: Hold at 95 °C for 30 minutes, then cool from 85 °C to 50 °C at -0.04 °C min$^{-1}$, then cool from 50 °C to room temperature at -0.5 °C min$^{-1}$.
- **Long anneal**: Hold at 95 °C for 30 minutes, then cool from 85 °C to 40 °C at -0.01 °C min$^{-1}$, then cool from 40 °C to room temperature at -0.1 °C min$^{-1}$.

Unless otherwise specified, the short anneal protocol was used for the preparation of unary C-star condensates, and the long anneal protocol for the preparation of binary C-star condensates. Condensates were extracted from capillaries by scoring each end with a diamond scribe, snapping at the scored lines, and placing the cut capillary vertically into an Eppendorf containing 0.3 M NaCl in TE for a minimum of 10 minutes. For large capillaries, 60 $\mu$L of buffer was used; for small capillaries, 30 $\mu$L. Fluorescent D1 nanostars, used in permeability experiments, were prepared to a final concentration of 10 $\mu$M in 0.3 M NaCl in TE by mixing stoichiometric ratios of all core strands — *i.e.* all component strands except the cholesterolised terminal strands — into an Eppendorf tube and annealing using the short anneal protocol.

#### Disassembly of C-star condensates

For condensate disassembly driven by toehold-mediated strand displacement, the appropriate invader strands (see Supplementary Table S2), reconstituted to 100 $\mu$M in 0.3 M NaCl in TE, were added to C-star condensates at a 6.6 $\times$ excess and allowed to incubate at room temperature. Invader strands were not actively mixed into the C-star condensate mixture, rather they were allowed to diffuse into the condensate, leading to a variation in disassembly times. For DNase I-mediated disassembly: DNase buffer (10 mM Tris-HCl, 2.5 mM $MgCl_2$, 0.5 mM $CaCl_2$, New England Biolabs) was used to dilute DNase I (RNase-free, New England Biolabs) at a 1:10 ratio. 15 $\mu$L of the enzyme in buffer was added to 7.5 $\mu$L of C-stars (equivalent to approximately 2.8 $\mu$g), and the activity of DNase on C-stars was monitored through confocal microscopy. All disassembly experiments were carried out at room temperature, and activity over time was monitored through confocal or epifluorescence microscopy.

#### Vesicle preparation and lipid shell formation

100 nm vesicles were prepared via extrusion — for all experimental variants, lipid films were prepared by evaporating ethanol-stabilised chloroform from the appropriate lipid mixture and drying under vaccuum overnight. Films were the rehydrated at 2 mg mL$^{-1}$ in buffer, freeze-thawed five times, and passed 21 times through 100 nm polycarbonate membranes (Whatman Nucleopore) using an Avanti Polar Lipids Mini-Extruder. Vesicle dimensions were verified through dynamic light scattering using a Malvern Panalytical Zetasizer, with typical data shown in Fig. S2. Experimental variants:

- Unlabelled SUVs: DOPC only; film rehydrated in 0.3 M sucrose in TE.
- Labelled SUVs: 0.8 mol% DHPE-Texas Red in DOPC; film rehydrated in 0.3 M sucrose in TE.
- Encapsulating calcein: DOPC only; film rehydrated in 50 mM calcein in 0.3 M sucrose in TE. Size Exclusion Chromatography with Sephadex G50 resin was used to separate calcein-loaded vesicles from unencapsulated calcein.
- Probing SUV movement on condensate surface: SUVs mixtures were prepared at a volume ratio of 1 labelled SUV to 799 unlabelled SUVs.

Extracted condensates were typically incubated with vesicles at a volume ratio of 10 to 1 for 10 minutes. Longer incubation times (up to 1 h) were found to have no significant effect on lipid shell properties (see Fig. S3).

#### Lipid shell permeability

Unlabelled vesicles were added to the extracted A1 C-star condensates at the ratio described above and incubated for 10 minutes to coat with a lipid shell. Aliquots of these were transferred to imaging wells, to which fluorescent probes were added. Probe concentrations used were as follows: 0.1 $\mu$M calcein; 1 $\mu$M 20 kDa TRITC-Dextran, 3 $\mu$M D1 nanostars, and 0.2 mg/mL Texas Red-labelled streptavidin. Total volume of condensate, probe, and 0.3 M NaCl in TE buffer in imaging wells was kept constant across all experiments. Samples were incubated with probe for 10 minutes prior to image acquisition. Three independent repeats of each probe variant were carried out, and between 7 and 25 (median of 12) condensates were imaged per repeat. Error bars show the standard error. Images were acquired with confocal microscopy, and custom code was used to segment images to extract and calculate a ratio of internal versus background fluorescence intensity. Control experiments were carried out with DOPC Giant Unilamellar Vesicles (GUVs) produced *via* electroformation in 0.3 M sucrose. Permeability experiments with a lipid shell formed from a longer incubation time (see Fig. S3) used an incubation time of 1 h. All other experimental parameters were kept constant. Two repeats were executed for each probe, with 11 to 13 lipid-coated condensates imaged per repeat.

#### Transcription of Broccoli in binary C-star condensates

A2-B1 binary condensates were prepared and extracted into 0.3 M NaCl in TE as previously described. To a 20 $\mu$L aliquot of condensates, 2 $\mu$L of vesicles were added and incubated for a maximum of 1 hour. This and a separate aliquot of non-lipid-coated condensates were washed three times using 0.3 M NaCl in TE. Following this, lipid-coated and non-coated condensates were loaded into separate wells containing 10 $\mu$L 0.3 M NaCl in TE at a target

C-star end concentration of 1.4 $\mu$M. B1 invader strands were added to these wells at $12\times$ excess for 30 min to ensure complete disassembly of B1 C-stars, after which all wells were washed three times with 0.3 M NaCl in TE. Etching was monitored with epifluorescence microscopy (see below) with image acquisition every 30 seconds for 30 minutes. For synthetic cells containing a fluorescently-labelled Bridge strand, Alexa Fluor 647 was excited at 640 nm. For synthetic cells coated in a fluorescently-labelled lipid shell, Texas Red-DHPE was excited at 590 nm. Expression of Broccoli aptamer was undertaken at room temperature. With the condensate extraction and washing steps described above, precise control over the number and size of condensates in imaging wells cannot be achieved. To mitigate instances where condensate concentration may exceed the targeted end concentration of 1.4 $\mu$m, we used transcription buffer, DTT, NTPs, and RNase inhibitor ratios of twice those described in the protocol for the CellScript T7-FlashScribe Transcription Kit.[2] Transcription was monitored using epifluorescence microscopy, imaging at regular intervals for a minimum of 18 hours (20 minutes for the first 2 hours, then 30 minutes until experiment end). Broccoli·DFHBI was excited at 470 nm, the Alexa Fluor 647-labelled Bridge strand was excited at 640 nm, and Texas Red-DHPE was excited at 590 nm. Synthetic cells containing the Alexa Fluor 647-labelled Bridge strand used a lipid shell made with unlabelled SUVs, whereas those with a Texas Red-labelled lipid shell used an unlabelled Bridge strand. Alexa Fluor 647 and Texas Red were not used in the same system. Experiments monitoring bulk transcription of (non-lipid-coated) condensates and free template (Fig. S7) were carried using the same protocol in well plates, using the unlabelled A2.Bridge strand, and monitoring fluorescence with a BMG CLARIOstar Plus plate reader set to excite wells at a 10 nm band centred at 447 nm and capture fluorescence emission between 496 nm and 506 nm.

#### Imaging techniques

Most confocal micrographs were acquired using a Leica TCS SP5 confocal microscope with an HC PL FLUOTAR $20\times$ 0.50 N.A. dry objective (Leica). Fluorescein-labelled samples were illuminated using Ar-ion laser lines (exciting fluorescein and calcein at 496 nm) or He-Ne laser lines (543 nm for TRITC, 594 nm for Texas Red). Micrographs monitoring for movement of SUVs on condensate surface (Fig. 2(c)) were obtained using a Leica STELLARIS 8 Inverted Confocal Microscope using an HC PL APO $20\times$ 0.75 N.A. CS2 dry objective (Leica), using a tunable white light laser to excite the Texas Red-labelled lipid with multiple wavelengths between 555 nm and 595 nm at 8 nm intervals.

Epifluorescence micrographs were obtained using a Nikon Eclipse Ti2-E inverted microscope, equipped with a digital camera (Hamamatsu ORCA-Flash4.0 V3), a tunable light source (Lumencor SPECTRA X LED engine), and Plan Fluor $20\times$ 0.75 N.A and Plan Fluor $40\times$ 0.9 5 N.A dry objectives (Nikon).

All samples were imaged in polydimethylsiloxane (PDMS) wells plasma bonded to glass cover slides, with the wells and cover slides cleaned in ethanol and thoroughly dried prior to plasma bonding. Wells were sealed with MicroAmp Optical Adhesive Film (ThermoFisher Scientific) to prevent evaporation during imaging.

#### Image analysis

Custom code was written for all image segmentation using MATLAB R2019b+, with the Image Processing and Statistics and Machine Learning toolboxes installed. Micrographs were exported as Tag Image File Format files automatically labelled by colour channel, timepoint and z-stack position (where relevant). Wherever possible, image segmentation was carried out on very isolated condensates, *i.e.* those at a reasonable distance from others. Brief descriptions of image processing steps for the various experiments are described below:

- **Monitoring C-star disassembly:** Confocal images were cropped to the region of interest. Masks of condensates were created by applying a Gaussian filter to the image channel containing the fluorophore of interest and thresholding to detect pixels above a certain fluorescence intensity to create a binary image. Mask area at each timepoint was extracted using the *regionprops* function. Data from this analysis are presented in

Fig. 1(g) and Fig. 2(e)*iv* and (g)*iii*.

- **Lipid shell permeability:** Confocal images were cropped to the region of interest. Image contrast was increased and Gaussian filtering applied, and these processed images were segmented using k-means clustering to create masks for the background and the inside of the condensate. The generated masks were then manually checked to ensure clusters were labelled correctly. To ensure edge effects did not influence results, both masks were eroded (*imerode*) using a disk-shaped structuring element with radius defined by the highest of either the diffraction limit (reduced Rayleigh criterion) or the calculated projection (using condensate size determined by mask area and assuming perfectly spherical). The refined masks were applied to raw confocal images to determine average fluorescence intensity inside and outside the condensates. Data from this analysis are presented in Fig. 2(d) and Fig. S3.

- **Extent of phase separation in binary condensates:** For confocal images of fluorescently-labelled binary condensates with a large field-of-view: first, Gaussian filtering was applied, after which the average background fluorescence intensity was determined through manual sampling. This value was subtracted from the filtered image, which was then binarised to create a mask containing all condensates in the image. The mask was refined with *regionprops* to include circular condensates (circularity $> 0.9$) of the desired size (diameter between $5\,\mu$m and $25\,\mu$m), to ensure condensates which have aggregated or are too small to accurately threshold are excluded from the analysis. Each individual condensate was then segmented using bespoke thresholding, yielding masks for the fluorophore- rich and fluorophore- poor phases which were then each eroded using the methods described previously to mitigate the impact of edge effects on the final calculation. The average fluorescent intensity of each phase of the binary condensate was then extracted and used to calculate an intensity ratio per condensate. This intensity ratio reflects the average fluorescence intensity of the inner (fluorophore-poor) phase divided the average intensity in the outer (fluorophore-rich) phase. This approach was used to analyse 164 A2-B1 condensates prepared without the BT duplex and 149 A2-B1 condensates prepared with the BT duplex, where the B1 motif is labelled with a fluorescein probe on the inner junction of the nanostar. Data from this analysis are presented in Fig. 3(c), plotted using open source code for generating violin plots, using a kernel density estimate bandwidth of 0.03.[3]

- **Size and local fluorescence intensity of synthetic cells transcribing Broccoli:** See Fig. S11 for workflow description supplemented with example images. Epifluorescence micrographs were cropped to the region of interest, containing the condensate and sufficient background area. Using bright-field images at the start and end of the transcription run for reference, a line was drawn spanning the condensate bounds and background. Where necessary, images were rotated to ensure the drawn line was straight and horizontal, and from this, line profiles of Alexa Fluor 647 fluorescence intensity were plotted from a 10 pixel average normal to the line (5 px above, 5 px below, to reduce noise). Peak detection and filtering based on prominence was applied to the gradient of the extracted line profile and the difference between the pixel position of outermost peaks was used to track condensate size. To track local fluorescence intensity, polygonal masks were hand-drawn to segment regions inside and outside the condensate. Hand-drawn masking was necessitated as semi-automated approaches led to inaccurate segmentation of the etched region inside the condensate, tending to include the non-etched A2-rich phases. Data from this analysis are presented in Fig. 4(e).

### Supplementary Tables

Table S1: Description and list of component strands of C-star and nanostar populations used in this work. For fluorescent labelling of populations, the appropriate strand was substituted with its fluorescent equivalent detailed in Table S2, which also lists the appropriate invader strand sequences for C-star disassembly *via* toehold-mediated strand displacement.

| Population name | Component strands | Description |
|---|---|---|
| A1 | A1.Core1; A1.Core2; A1.Core3; A1.Core4; CholTerminal | C-star with arm length of 35 bp |
| A2 | A2.Core1; A2.Core2; A2.Core3; A2.Core4; A2.Base; A2.Bridge; A2.Template; CholTerminal | Broccoli-templating C-star, arm length 35 bp |
| B1 | B.Core1; B.Core2; B.Core3; B.Core4; B1.Bridge; B1.TerminalComplement; B.CholTerminal | TMSD-modified C-star, arm length 50 bp |
| B2 | B.Core1; B.Core2; B.Core3; B.Core4; B2.Bridge; B2.TerminalComplement; B.CholTerminal | TMSD-modified C-star, minor variation on B1 design, arm length 50 bp |
| B3 | B.Core1; B.Core2; B.Core3; B.Core4; B3.Bridge; B3.TerminalComplement; B.CholTerminal | TMSD-modified C-star, minor variation on B1 design, arm length 50 bp |
| C1 | C1.Core1; C1.Core2; C1.Core3; C1.Core4; C1.Bridge; C1.TerminalComplement; C.CholTerminal | TMSD-modified C-star, arm length 48 bp |
| D1 | D1.Core1; D1.Core2; D1.Core3; D1.Core4; CholTerminal | C-star with arm length of 28 bp |
| D1 (nanostar) | D1.Core1; D1.Core2; D1.Core3; D1.Core4 | Nanostar with arm length of 28 bp |

Table S2: Oligonucleotide sequences for all materials used in this work, written in 5' to 3' direction and split into groups of three for clarity.

| Strand name | Oligonucleotide sequence |
|---|---|
| A1.Core1 | CGA CGC CGT GAC GCG TTG ATG ACT CGA CTG ACC AGA GCA TCT TAG CTC ACT GGA AAC |
| FluorA1.Core1 | CGA CGC CGT GAC GCG TTG ATG ACT CGA CTG ACC AG \Cy5\GCA TCT TAG CTC ACT GGA AAC |
| A1.Core2 | CGA CGC CGT GAC GCG TTT CCA GTG AGC TAA GAT GCA CGA ATG ACT GCA CTG TCA AAC |
| A1.Core3 | CGA CGC CGT GAC GCG TTT GAC AGT GCA GTC ATT CGA CGA ATC GAA ATA CTG TGG AAC |
| A1.Core4 | CGA CGC CGT GAC GCG TTC CAC AGT ATT TCG ATT CGA CTG GTC AGT CGA GTC ATC AAC |
| A2.Core 1 | CGA CGC CGT GAC GCC GTG GCC TGT GAT TGA GGC GCT GCG TCG TCC ACC GTG TGA AAC TTG TCC GTT CTA AAT C |
| A2.Core 2 | CGA CGC CGT GAC GCG TTT CAC ACG GTG GAC GAC GCT CGG ACT AGA ACT GTC TCG AAC |
| A2.Core 3 | CGA CGC CGT GAC GCG TTC GAG ACA GTT CTA GTC CGT CGC GAA TAC GCC GTG CCG TGC |
| A2.Core 4 | CGA CGC CGT GAC GCG CAC GGC ACG GCG TAT TCG CGT GCG CCT CAA TCA CAG GCC ACG |
| A2.Base | GGT GAG GTG AGT GGG ATT TAG AAC GGA C |
| A2.Bridge | CCA CTC ACC TCA CCT AAT ACG ACT CAC TAT A |

*Continued on next page*

Table S2: Oligonucleotide sequences for all materials used in this work, written in 5' to 3' direction and split into groups of three for clarity.

| Strand name | Oligonucleotide sequence |
|---|---|
| FluorA2.Bridge | \Alexa647\CCA CTC ACC TCA CCT AAT ACG ACT CAC TAT A |
| A2.Template | GGG TCT AGG AGC CCA CAC TCT ACT CGA CAG ATA CGA ATA TCT GGA CCC GAC CGT CTC CTA GAC CCT ATA GTG AGT CGT ATT A |
| B.Core1 | TTC CCT GGC GGC GAT TCT CGA TCA GCG CTA ATC A |
| FluorB.Core1 | TTC CCT GGC GGC GAT TCT CGA \Fluorescein-dT\CAG CGC TAA TCA |
| B.Core2 | TTC CCT GGC GGA ACC TAT ACC TTC GAG AAT CGC C |
| B.Core3 | TTC CCT GGC TAT GCT GCT CTG TGG TAT AGG TTC C |
| B.Core4 | TTG GGT GGC TGA TTA GCG CTG TCA GAG CAG CAT A |
| B1.Bridge | CAT CTT GCC AGG GAA GTG AAG AAA CTG TCC |
| B1.TerminalComplement | ACT ACA CGT CTC AGG GAC AGT TTC TTC AC |
| B1.Invader | GGA CAG TTT CTT CAC TTC CCT GGC AAG ATG |
| B2.Bridge | CAT CTT GCC AGG GAA TTA CCG ACC CTT TCC |
| B2.TerminalComplement | ACT ACA CGT CTC AGG GAA AGG GTC GGT AA |
| B2.Invader | GGA AAG GGT CGG TAA TTC CCT GGC AAG ATG |
| B3.Bridge | CAT CTT GCC AGG GAA TTA CCG ACT CAT GAC |
| B3.TerminalComplement | ACT ACA CGT CTC AGG TCA TGA GTC GGT AA |

*Continued on next page*

Table S2: Oligonucleotide sequences for all materials used in this work, written in 5' to 3' direction and split into groups of three for clarity.

| Strand name | Oligonucleotide sequence |
|---|---|
| B3.Invader | GTC ATG AGT CGG TAA TTC CCT GGC AAG ATG |
| C.Core1 | CCT ACT TCC ACC GCG TCG CAG CAC TGC CCG TCC TCG C |
| FluorC.Core1 (Fluor) | CCT ACT TCC ACC GCG TCG CAG CAC \Fluorescein-dT\GC CCG TCC TCG C |
| C1.Core2 | CCT ACT TCC ACC GCG AGG ACG GGC TCG TCG CTT TCG C |
| C1.Core3 | CCT ACT TCC ACC GCG AAA GCG ACG TCG TCA ACC GAA C |
| C1.Core4 | CCT ACT TCC ACC GTT CGG TTG ACG TGT GCT GCG ACG C |
| C1.Bridge | GTG TGA GGT GGA AGT AGG GTG AGG TGA TGG |
| C1.TerminalComplement | GCG GAG GTC ACG CCA TCA CCT CAC |
| C1.Invader | CCA TCA CCT CAC CCT ACT TCC ACC TCA CAC |
| D1.Core1 | CGA CGC CGT GAC GCG CTT GGG CGT GGC GTC GCG AGC GCC AAG C |
| FluorD1.Core1 | CGA CGC CGT GAC GCG CTT GGG CGT GGC G\Fluorescein-dT\C GCG AGC GCC AAG C |
| D1.Core2 | CGA CGC CGT GAC GCG CTT GGC GCT CGC GTC GGC CAT TGA CTG C |
| D1.Core3 | CGA CGC CGT GAC GCG CAG TCA ATG GCC GTC GGC CAC GCG CAC G |

*Continued on next page*

Table S2: Oligonucleotide sequences for all materials used in this work, written in 5' to 3' direction and split into groups of three for clarity.

| Strand name | Oligonucleotide sequence |
|---|---|
| D1.Core4 | CGA CGC CGT GAC GCC GTG CGC GTG GCC GTC GCC ACG CCC AAG C |
| CholTerminal | GCG TCA CGG CGT CGA A \TEG-Cholesterol\ |
| B.CholTerminal | CTG AGA CGT GTA GT \TEG-Cholesterol\ |
| C.CholTerminal | CGT GAC CTC CGC AA \TEG-Cholesterol\ |

### Supporting Figures

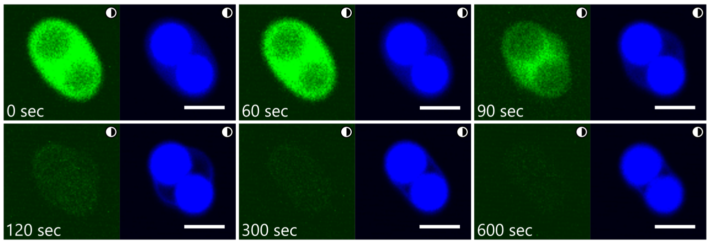


Figure S1: **Targeted disassembly of binary condensates reveals incomplete demixing.** Confocal micrographs showing the targeted disassembly (*via* TMSD) of B2 C-stars in the A1-B2 binary condensate shown in Fig. 1(f), where image contrast has been increased to show otherwise very faint detail. Leftmost image in each panel shows the fluorescence channel of the fluorescein-labelled B2 condensates (green), rightmost images show the fluorescence channel of the Cy5-labelled A1 condensates, coloured blue. By greatly increasing image contrast, we can more easily observe a faint halo in the Cy-5 (blue) channel caused by a low-density mesh of A1 C-stars trapped in the outer phase of the binary condensate (at 120 seconds). This mesh collapses onto the A1-rich cores over time. Timestamps mark time elapsed after adding the invader strand. Scale bars 10 $\mu$m.

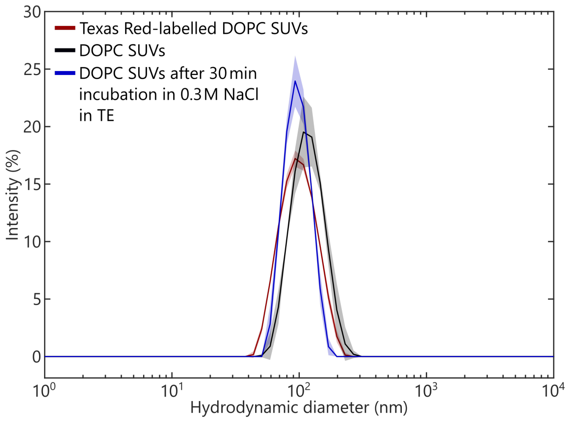


Figure S2: **Dynamic light scattering (DLS) confirms SUV size.** Intensity distribution of the hydrodynamic diameter of small unilamellar vesicles (SUVs) as determined with DLS. Data are relative to typical SUV batches, formed *via* extrusion in TE buffer supplemented with 0.3 M sucrose. The red line shows data for DOPC SUVs stained with Texas Red-labelled DHPE, the black line shows DOPC-only SUVs, and the blue line shows DOPC-only SUVs after 30 minutes of incubation in the buffer used for annealing and extracting C-star condensates (0.3 M NaCl in TE). Despite the osmolarity difference, DLS data indicates no significant change in the size of SUVs after incubation in 0.3 M NaCl in TE. Solid lines show the mean of three measurements, with shaded regions indicating standard deviation.

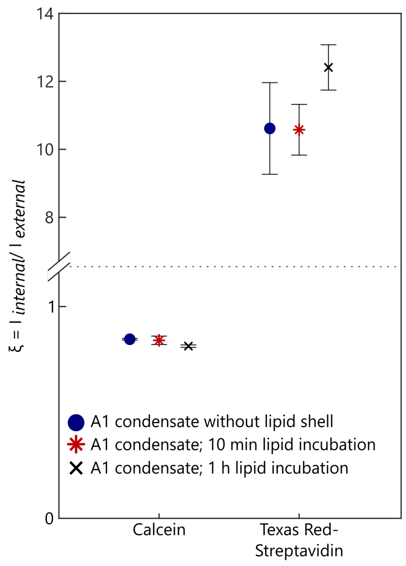


Figure S3: **SUV incubation time does not reduce permeability of lipid shells.** Partitioning of calcein and Texas Red-labelled streptavidin into A1 C-star condensates without a lipid shell (blue circles) and with lipid shells formed from an SUV incubation time of either 10 minutes (red stars) or 1 hour (black crosses), gauged using the ratio $\xi$ of probe fluorescence intensity inside ($I_{\text{internal}}$) and outside the condensate ($I_{\text{external}}$), measured from confocal micrographs. Data shows that lipid shells formed from an increased SUV incubation time remain permeable to the tested probes. For condensates without and with a lipid shell from 10 minute incubation, three independent repeats were executed, sampling between 7 and 19 (median of 13) condensates per repeat. For condensates with a 1 hour SUV incubation time, two independent repeats were executed, sampling between 11 and 13 condensates per repeat. Error bars show the standard error.

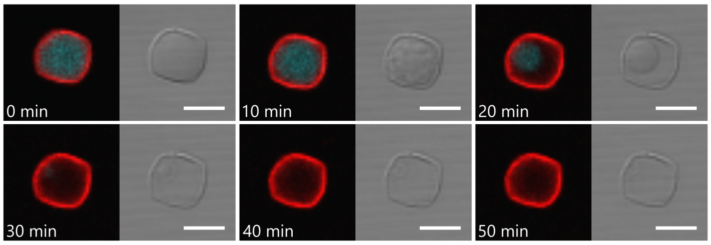


Figure S4: **Enzyme-triggered disassembly of lipid-coated DNA condensates.** DNase I disassembly of a D1 C-star condensate (fluorescein-labelled, arm length 28 bp) enveloped in a Texas Red-labelled DOPC lipid shell. DNase I digests the condensate, leaving behind the lipid shell. Timestamps mark time elapsed after DNase I addition. Scale bars 20 $\mu$m.

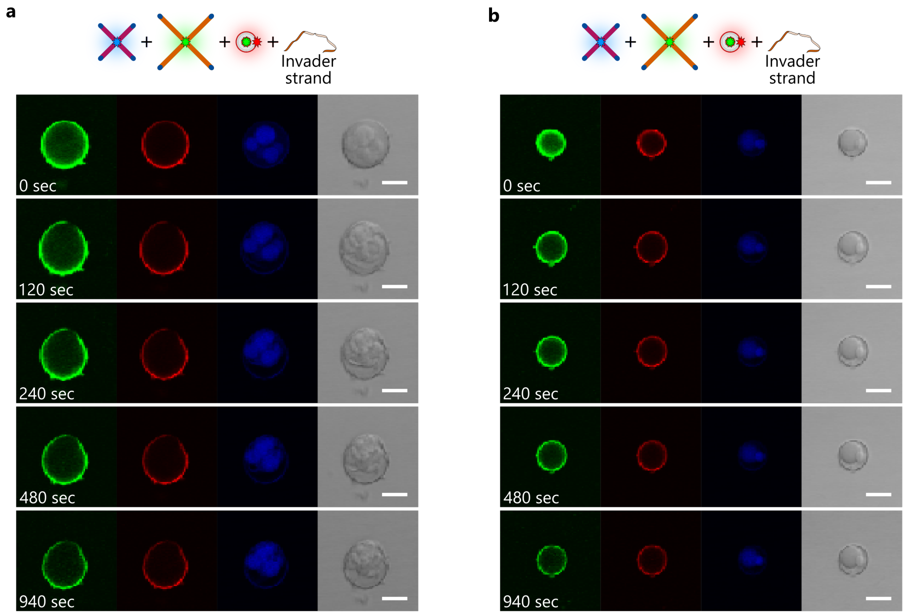


Figure S5: **Etching of binary C-star condensates with a lipid shell composed of calcein-loaded, Texas Red-labelled SUVs.** Confocal and bright-field images of two A1-B3 binary C-star condensates undergoing etching of the B3 motif through toehold-mediated strand displacement. In both panels **(a)** and **(b)**, imaging channels are, from left to right: green fluorescence channel showing the signal from calcein loaded in the SUVs and fluorescein-labelled B3 motifs in the outer shell of the condensates; red fluorescent channel showing Texas Red-DHPE labelling the SUV membranes; blue fluorescent channel showing the Cy5-labelled A1 motifs in the core of the condensates; bright-field channel. While the signal in the green channel is dominated by the calcein, etching of the outer phase can be noted from darkening of the region within the lipid shell, as well as from the bright-field images. SUVs are found to retain calcein after B3 disassembly. Scale bars 20 $\mu$m.

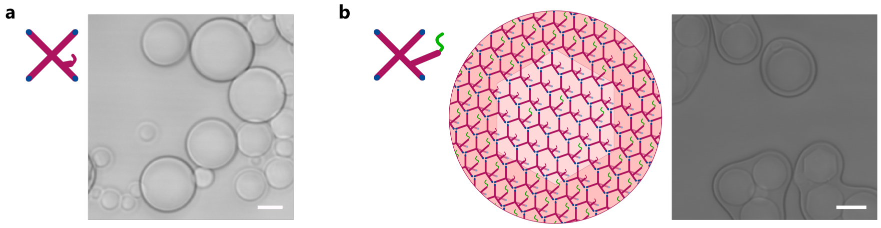


Figure S6: **Introduction of bulky Bridge-Template motifs induces de-mixing in single-component C-star condensates.** **(a)** Simplified schematic and bright-field micrograph of A2 condensates (arm length 35 bp) annealed without the Bridge and Template strands, showing no internal phase separation. **(b)** Schematic and bright-field micrograph of A2 condensates annealed with the Bridge and Template strands and exhibiting distinct de-mixing. All scale bars 10 $\mu$m.

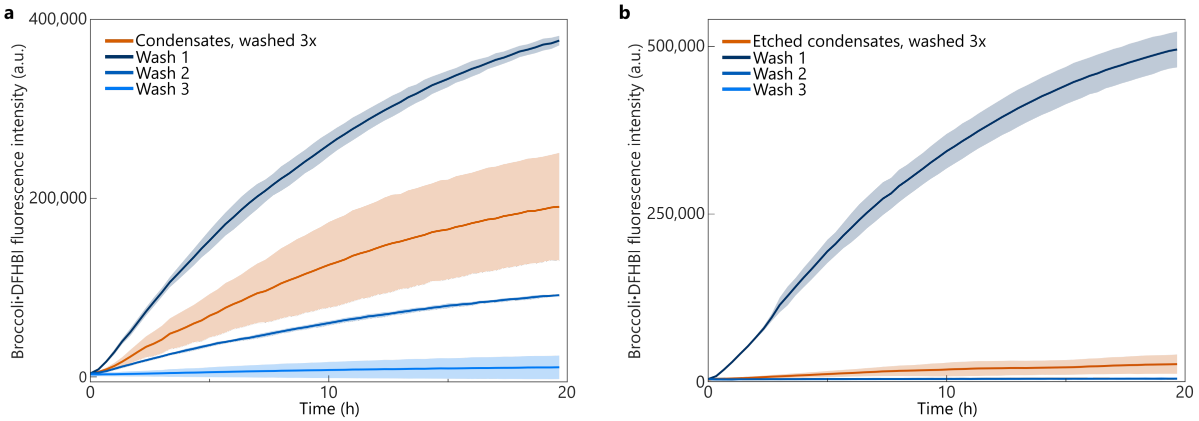


Figure S7: **Excess DNA template is removed by washing condensates, and targeted phase etching of binary condensates leads to the release of significant quantities of template.** Fluorimetry data recorded from transcription reactions producing the Broccoli aptamer from templates embedded in condensates, or present in the supernatants collected after condensate washing. **(a)** A2-B1 condensates are washed three times (see Methods). Transcription reactions carried out with the thrice-washed condensates and an equal volumes of the supernatants removed at each washing step reveal that a significant quantity of Template is initially present in the bulk (Wash 1), but can be removed with subsequent washing steps. **(b)** B1 C-stars in the thrice washed A2-B1 condensates are disassembled *via* toehold-mediated strand displacement (see Methods), and condensates are subsequently washed three times. The strong signal recorded from the supernatants collected during the first washing step after disassembly (Wash 1) indicates that a significant amount of template is incorporated into the B1-rich phase and released in the bulk upon disassembly. For all panels, data are presented as a mean (solid lines) $\pm$ the standard deviation (shaded regions) of three repeats per sample.

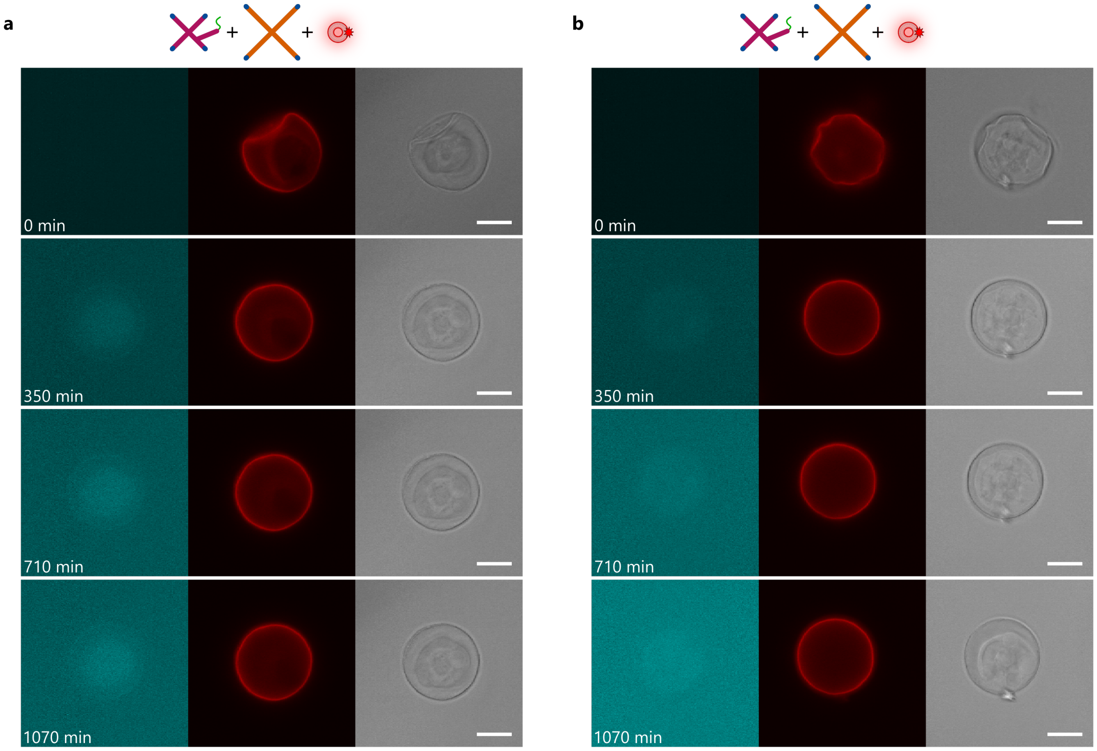


Figure S8: **Morphological evolution of synthetic cells with a fluroescently-labelled lipid shell during RNA transcription.** Both panels show epifluorescence (left and middle) and bright-field (right) micrographs of an RNA-transcription reaction sustained by A2-B1 synthetic cells. Leftmost image shows the increase in Broccoli fluorescent signal as transcription proceeds. These synthetic cells were coated with a Texas Red-labelled lipid shell (middle image, shown in red) and etched to disassemble B1 C-stars prior to transcription. We observe the same morphological response with these synthetic cells as in Fig. 4, Fig. S9 and Fig. S10, and we further note that the lipid shell, made visible with Texas Red labelling, is unaffected by the transcription reaction conditions. Panel **(a)** shows the same synthetic cell as in Fig. 4(d) with more timepoints to demonstrate the evolution of the system. Scale bars 20 $\mu$m. Timestamps mark time elapsed after addition of T7 RNA polymerase and the transcription mixture.

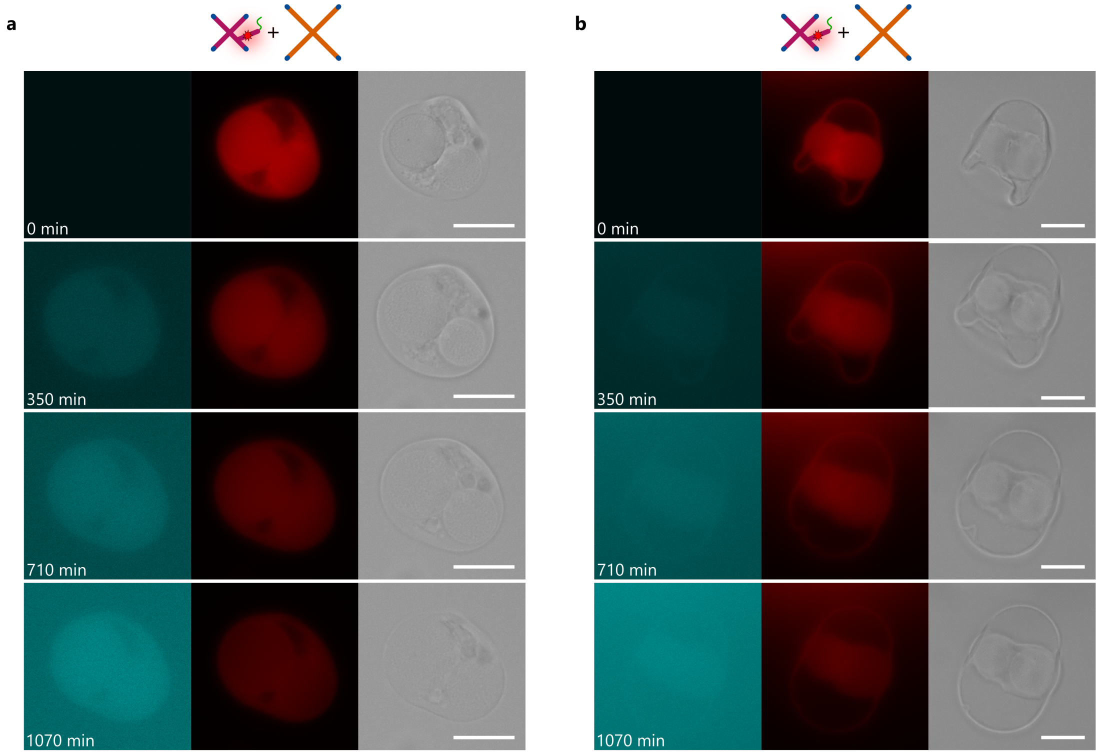


Figure S9: **Morphology of non-lipid-coated synthetic cells after etching and during transcription.** Both panels show epifluorescence (left and middle) and bright-field (right) micrographs of the RNA-templating A2-B1 binary synthetic cells undergoing transcription, after having been etched to disassemble the B1 motif. In this system, the Bridge strand in the A2 motif is labelled with Alexa Fluor 647 (middle image, shown in red). Leftmost images in each panel show the Broccoli fluorescent signal in cyan. The synthetic cells are not coated with a lipid shell. In roughly half of the analysed constructs, the synthetic cells adopt a compact morphology akin to that shown in panel **(a)**. Panel **(b)** shows the typical morphology of the other half of the constructs, where the A2-rich dense cores in the synthetic cell are encircled by a thin ring of A2-rich material (identified *via* fluorescent labelling of the A2 Bridge strand). Scale bars 20 $\mu$m. Timestamps mark time elapsed after addition of T7 RNA polymerase and the transcription mixture.

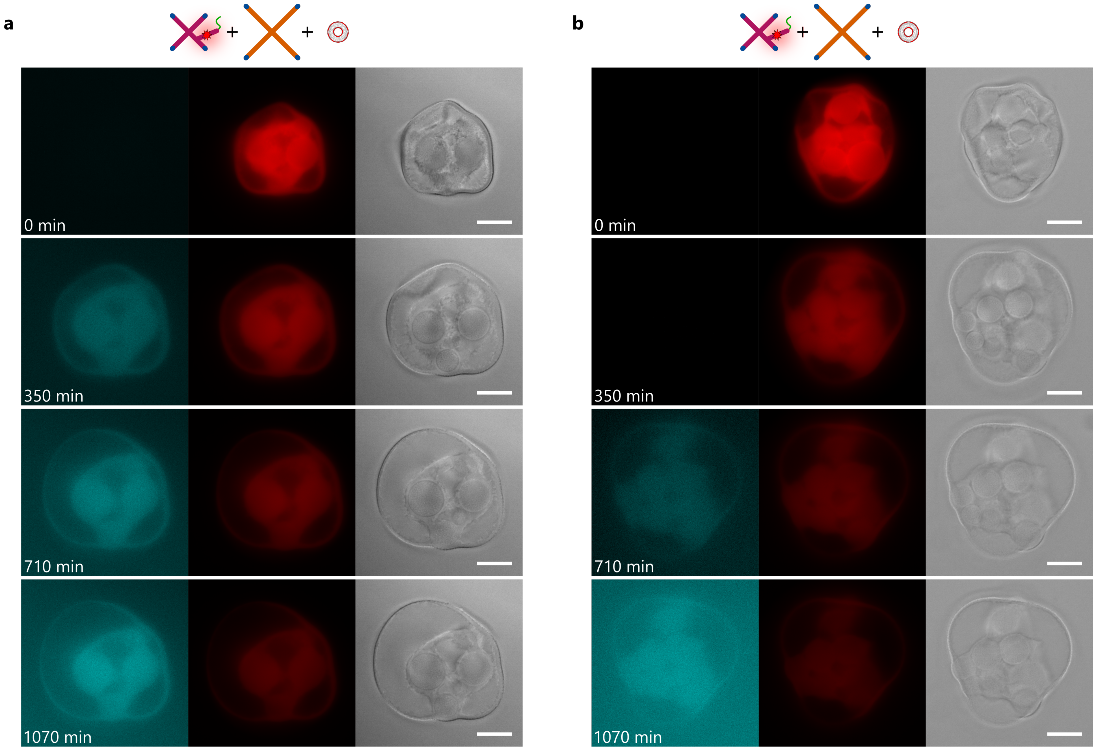


Figure S10: **Transcription-induced morphological response in lipid-coated synthetic cells.** Panels **(a)** and **(b)** show epifluorescence (left and middle) and bright-field (right) micrographs of an RNA-transcription reaction sustained by two different A2-B1 synthetic cells which were etched to disassemble B1 C-stars (as in Fig. 4), where the Bridge strand in the A2 motif is labelled with Alexa Fluor 647 (middle, shown in red). As transcription proceeds, the Broccoli fluorescent signal, shown in cyan in the leftmost image, increases, and the swelling response discussed in the main text is observed. Scale bars 20 $\mu$m. Timestamps mark time elapsed after addition of T7 RNA polymerase and the transcription mixture.

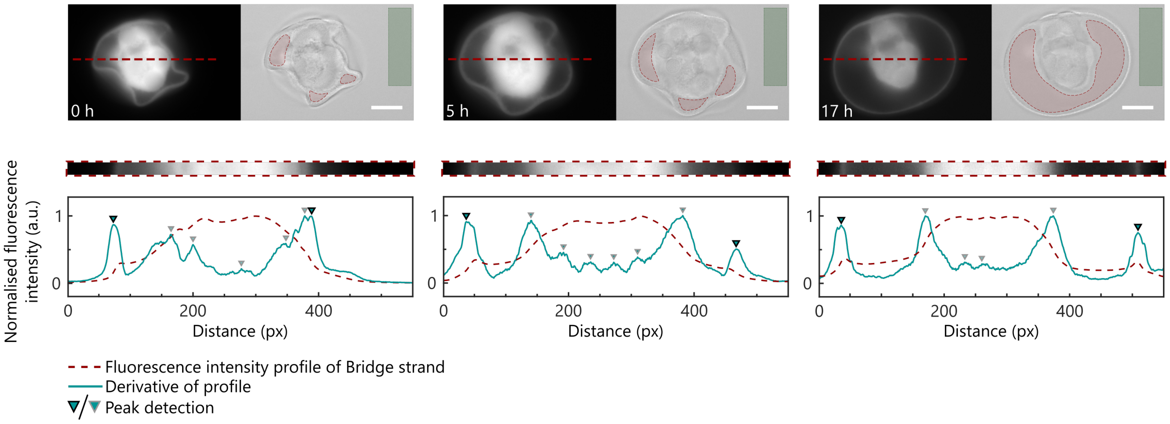


Figure S11: **Segmentation of a typical etched A2-B1 binary condensate to determine size and evaluate fluorescence intensity.** Fluorescence micrographs show the signal from the Alexa Fluor 647-labelled Bridge strand. A line is drawn on the condensate and applied to the Alexa Fluor 647 channel, from which an intensity profile is generated using an average of 5 px above and below the red dotted line (slice shown below micrographs). Condensate size is extracted by finding the outermost peaks in the gradient (solid cyan line) of the fluorescence intensity profile (dashed red line). Overlaid on the bright-field micrographs are hand-drawn masks showing regions inside (red shading) and outside (green shading) the condensate's bounds, taking care to avoid the non-etched A2-rich phase. These masks are applied to micrographs of the Broccoli·DFHBI fluorescence channel (not shown) and used to calculate the fluorescence intensity ratio $\zeta$ presented in Fig. 4(e)$i$. Scale bars 20 $\mu$m.

### List of Supporting Videos

- **Video V1:** Confocal and bright-field z-stack of non-lipid-coated binary A1-B3 condensates, where A1 C-stars are labelled with Cy5 (shown in blue) and B3 C-stars are labelled with fluorescein (shown in green).

- **Video V2:** Confocal and bright-field timelapse showing TMSD-mediated etching of the condensates detailed in Supporting Video V1 following addition of an invader strand designed to disassemble the B3 C-star motif.

- **Video V3:** Confocal and bright-field z-stack of a Texas Red-labelled lipid shell (shown in red, right) which remain after disassembly of a fluorescein-labelled B1 condensate (shown in green, middle). Contrast of the fluorescein channel has been enhanced for visibility.

- **Video V4:** Confocal and bright-field z-stack of a binary D1-C1 condensate coated in a lipid shell. D1 C-stars are labelled with fluorescein and shown in cyan, C1 C-stars are unlabelled, and the lipid shell is labelled with Texas Red, shown in red.

- **Videos V5 - V7:** Bright-field and epifluorescence timelapses showing the transcription of lipid-coated, etched, binary RNA-templating A2-B1 C-star condensates. The leftmost channel shows the signal from Broccoli·DFHBI fluorescence (shown in cyan), and the middle channel shows signal from the Texas Red-labelled lipid shell (shown in red). The A2.Bridge strand used in this system is non-fluorescent.

- **Videos V8 - V10:** Bright-field and epifluorescence timelapses showing the transcription of lipid-coated, etched, binary RNA-templating A2-B1 C-star condensates. The leftmost channel shows the signal from Broccoli·DFHBI fluorescence (shown in cyan), and the middle channel shows signal from the Alexa Fluor-labelled A2.Bridge strand (shown in red). The lipid shell is not labelled with a fluorophore.

- **Videos V11 - V13:** Bright-field and epifluorescence timelapses showing the transcription of etched, binary RNA-templating A2-B1 C-star condensates. The condensates are not coated in a lipid shell. The leftmost channel shows the signal from Broccoli·DFHBI fluorescence (shown in cyan), and the middle channel shows signal from the Alexa Fluor-labelled A2.Bridge strand (shown in red).
